# Gut microbiota assemblages of generalist predators are driven by local- and landscape-scale factors

**DOI:** 10.1101/2022.10.27.513979

**Authors:** Hafiz Sohaib Ahmed Saqib, Linyang Sun, Gabor Pozsgai, Pingping Liang, Mohsan Ullah Goraya, Komivi Senyo Akutse, Minsheng You, Geoff M. Gurr, Shijun You

**Affiliations:** State Key Laboratory for Ecological Pest Control of Fujian and Taiwan Crops, Institute of Applied Ecology, Fujian Agriculture and Forestry University, Fuzhou 350002, China; Guangdong Provincial Key Laboratory of Marine Biology, College of Science, Shantou University, Shantou 515063, China; Joint International Research Laboratory of Ecological Pest Control, Ministry of Education, Fuzhou 350002, China; Ministerial and Provincial Joint Innovation Centre for Safety Production of Cross-Strait Crops, Fujian Agriculture and Forestry University, Fuzhou 350002, China; Azorean Biodiversity Group, Centre for Ecology, Evolution and Environmental Changes, University of Azores, Portugal; Center for Infection and Immunity, Guangdong Provincial Engineering Research Center of Molecular Imaging, Guangdong Provincial Key Laboratory of Biomedical Imaging, the Fifth Affiliated Hospital, Sun Yat-sen University, Zhuhai, Guangdong, 519000, China; Guangdong Provincial Key Laboratory of Infectious Diseases and Molecular Immunopathology, Shantou University Medical College, Shantou, 515041, China; Plant Health Theme, International Centre of Insect Physiology and Ecology, Nairobi P.O. Box 30772-00100, Kenya; Gulbali Institute, Charles Sturt University, Orange, NSW 2800, Australia

**Keywords:** agroecosystem, microbiome, high throughput sequencing, microbe-environment interactions, Lycosidae

## Abstract

The gut microbiomes of arthropods are reported to have significant impact on key physiological functions such as nutrition, reproduction, behavior, and health. Spiders are diverse and numerically dominant predators in crop fields where they are potentially important regulators of pests. The taxonomic structure of spider gut microbiomes, and environmental drivers of composition are unknown. Harnessing spiders to support agricultural productivity is likely to be supported by an understanding of the gut microbiomes of these predators. This study aimed to deciphering the gut microbiome assembly of predators as well as elucidating the potential implications of key environmental constraints in this process. Here, we used high-throughput sequencing to examine for the first time how the assemblages of bacteria in the gut of spiders are shaped by diverse environmental variables. A total of 27 bacterial phyla were detected with Proteobacteria and Firmicutes dominant. The core bacterial communities included the families Enterobacteriaceae, Chloroplast, Lactobacillaceae, Pseudomonadaceae, Lachnospiraceae, Leuconostocaceae and Ruminococcaceae. Local drivers of microbiome composition were the globally-relevant input use system (organic production versus conventional practice), and crop identity (Chinese cabbage versus cauliflower). Landscape-scale factors, proportion of forest and grassland, compositional diversity, and habitat edge density, also strongly affected gut microbiota. Specific bacterial taxa were enriched in the gut of spiders sampled from different settings and seasons. These findings provide a comprehensive insight into the composition and plasticity of spider gut microbiota. Understanding the temporal responses of specific microbiota could lead to innovative strategies development for boosting biological control services of predators.

## 1 Introduction

Modern DNA-based methods have revealed that the guts of arthropods harbor a wide variety of microbes exerting strong effects on host fitness, including development, reproduction, host nutrition, stress tolerance against biotic and abiotic stresses, and regulation of host-pathogen interactions (Engel and Moran, 2013; Jang and Kikuchi, 2020). Research on bacterial diversity within arthropods recently focused on microbial communities’ interactions with their hosts. Studies have shown that symbiotic microbes within the host can be acquired from the external environment or through vertical transmission from other organisms (Hauke and Paul, 2011; Kwong et al., 2022). Therefore, understanding the environmental factors that determine the composition of gut microbes may thus provide insights into ecological interactions, particularly in highly dynamic ecosystems with continuously changing environmental conditions.

Gut microbial communities of arthropods are affected by the host diet (Marinozzi et al., 2013). Spiders are one of the most abundant and diverse groups of predators in agricultural fields, and exhibit a diversity of foraging, hunting, morphological and physiological traits that allow them to coexist (Viera and Gonzaga, 2017). These substantial evolutionary adaptations contribute to niche differentiation (Michalko et al., 2016) including utilisation of diverse prey. These factors allow spiders to persist in and exploit a wide range of spatio-temporal habitats including in-crop residency whilst utilizing non-pest prey prior to ‘switching’ when crop pests become available (Rand et al., 2006). The effects of these diet-related factors on the identity and composition of spider gut microbiota are likely to be significant but are largely unknown.

The gut microbiota of arthropods also varies geographically (Krawczyk et al., 2022), suggesting a role of factors in the surrounding environment such as microbes in locally available diet or foraging substrates. Spiders utilise a wide variety of prey located in contrasting habitat types such as foliage or the soil surface and, since the surrounding landscape composition drives the abundance and diversity of these food items (Saqib et al., 2021), it is plausible that spiders’ microbial communities can be affected by such factors. Effects on gut microbial diversity of arthropods from the surrounding habitat have been reported (Tiede et al., 2017). However, the effects of larger, landscape-scale effects, along with local factors such as farming practices, remain largely unclear.

Predator and prey species coexist in an environment full of toxins (such as plant defence compounds, pollutants and pesticides). Pesticides are widely used in conventional farming systems, which results in evolutionary adaptations in both predators and prey to counteract their effects (e.g., the mechanism of detoxification, changes in the target site, neutralization of toxins) (Köhler and Triebskorn, 2013). Although it is believed that all these resistance mechanisms are encoded within the insect genomes, omics analyses have revealed that a variety of organisms possess gut microbes, that actively degrade toxins including plant defence and pesticidal compounds (Itoh et al., 2018). Organic farming provides an alternative to conventional farming systems by utilizing crop rotation, pathogen-resistant cultivars, and organic fertilizers instead of synthetic chemicals. However, the effects of organic agriculture on biodiversity remain the subject of much study and are poorly understood in relation to gut microbiota.

In brassica vegetable growing systems, different species of *Brassica* are often cultivated in close proximity and such polycultures can affect the diversity of prey species available to predators (Brandmeier et al., 2021; N’Woueni and Gaoue, 2022). Furthermore, the prey assemblages vary with the changing cropping patterns and climate of different seasons (Liu et al., 2018; Radzikowski et al., 2020). Since the composition of the predator gut microbiome may be affected by diet, including potential acquisition of microbes from prey, a range of environmental drives is likely influence the spider microbiomes.

Few studies have been reported of the gut microbes of spiders (Kumar et al., 2020; Tyagi et al., 2021; Vanthournout and Hendrickx, 2015; Zhang et al., 2018, 2017), despite the foregoing range of factors that are likely to be drivers of spider performance as control agents of pests in crop systems. Accordingly, we employed 16S rDNA high-throughput sequencing to characterise the gut microbial assemblages of spiders sampled from brassica vegetable fields and evaluate the effects of local- and landscape-scale variables on taxonomic composition. Specially we hypothesized that the gut microbiome is highly plastic rather than fixed, with season, local agronomic factors and the wider landscape significantly influencing the assemblages of microbes in the gut of spiders.

## 2 Material and methods

### 2.1 Study system, area and design

In 2019, we sampled spiders from vegetable farms in Fujian Province, southeastern China, once every season for the four seasons (i.e., four times) at crop maturity (Fig. 1a). In this region, farms are typically smallholdings with highly dynamic, polyculture vegetable production systems. A total of 18 commercial crop fields were chosen, each at least 1 km apart, to represent various management systems and crop types, as well as varying fractions of land usage in the surrounding landscapes (Fig. 1b).

**Fig. 1.**
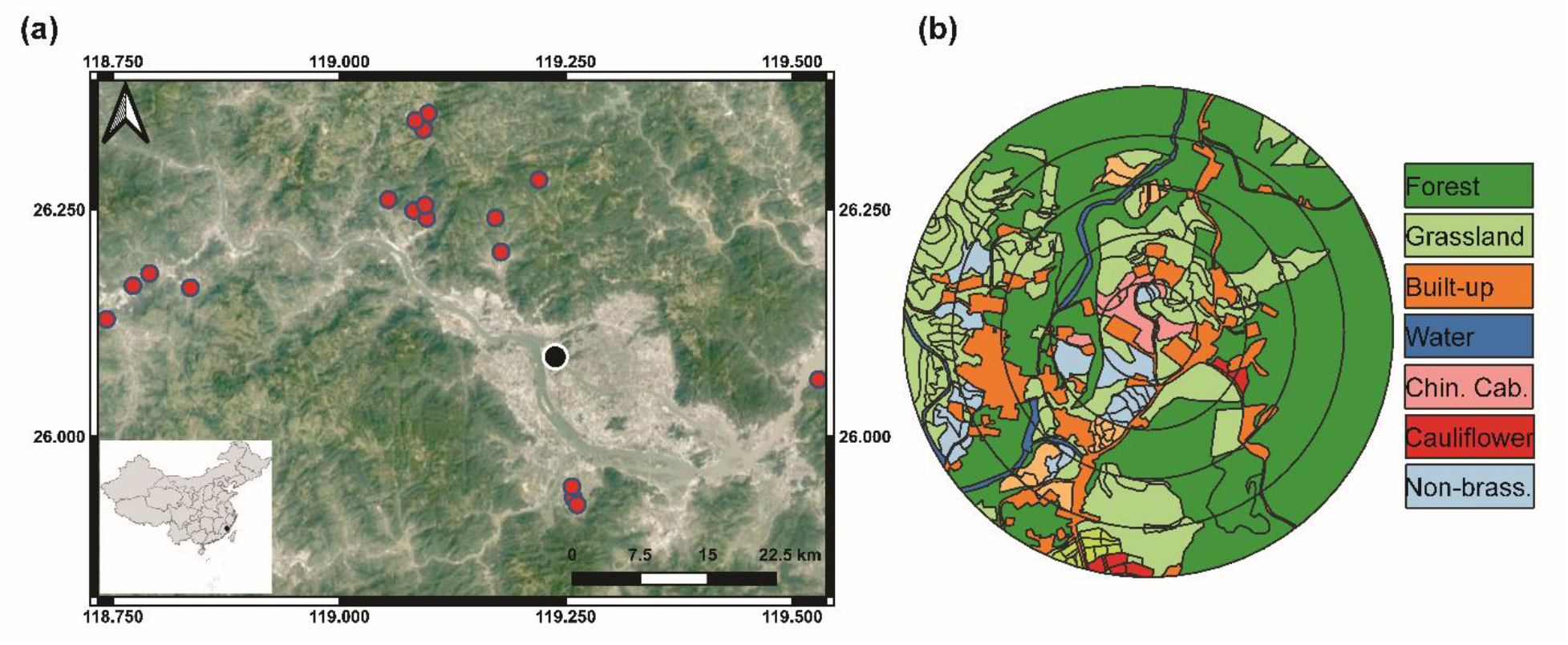
**(a)** Map showing the sampling locations in Fujian Province, southeastern China. **(b)** Example of landscape mapping of different habitats within 100, 200, 300, 400 and 500 m radius buffers around the focal sampling field.

Fields were divided into two groups based on management practices: 12 conventionally managed (i.e., synthetic pesticides or fertilizers were used) and six organically managed (i.e., no synthetic pesticides or fertilizers were used). Chinese cabbage (*Brassica rapa pekinensis*) and cauliflower (*Brassica oleracea*) were grown using direct sowing and seedling transplanting methods. Since samples were collected from farmers’ fields, we did not interfere or control the inputs in either conventional or organic farms nor did we intervene in any management practices. The disparity in field count between these two farming groups reflected their relative representation in this region. In all the four seasons, statistically adequate replicates of Chinese cabbage and cauliflower were represented in both organic and conventionally managed fields.

To ensure that the crops were not damaged and prevent surface DNA contamination, spiders were hand-collected using sampling method 4 described of Sørensen et al. (2002). Only spiders belonging to the Lycosidae family were collected since these were numerically dominant and known to be potentially important predators of brassica pests (ref). Individual lycosids were sampled from the soil surface and plants directly into 5 mL clean tubes, by two people for an hour of active searching per site. Samples were immediately transported to the laboratory in an icebox, and then kept at −80 °C.

### 2.2 Landscape analyses

A drone (PHANTOM 4, Shenzhen Dajiang Baiwang Technology Co., Ltd., China) was used to take aerial photographs of each field and the landscape to a 500 m radius circle to investigate the distribution of habitats. Aerial images were used to classify the vegetation types within the 500 m circles into grassland, forest, built-up (e.g., residential land, greenhouses, and roads), water surfaces (e.g., streams and ponds), Chinese cabbage, cauliflower, other Brassica crops (e.g., broccoli, canola, and mustard), non-brassica crops (e.g., pepper, eggplant, corn, and beans) and fallow land (arable having no crop) (Fig. 1b). The compositional diversity of landscape was calculated using Shannon-Wiener diversity index (SHDI). QGIS 3.4 was used to calculate the proportions of various habitat types and edge densities in the 500-meter radius landscape surrounding the focal field, which was divided into five concentric buffer circles at intervals of 100-meters.

### 2.3 DNA extraction, PCR and library preparation

A modified salt DNA extraction protocol (Sunnucks and Hales, 1996) was used to extract the genomic DNA of 732 adult spiders. Before performing DNA extractions, individual spiders were surface sterilized with 70% ethanol and washed three times with double distilled sterile water. Three lycosid individuals collected from the same field were pooled to perform a single DNA extraction. All the genomic DNA was kept at −80 °C until it was needed. Using the bacterial V3 – V4 barcode region, 16S ribosomal RNA (rRNA) was amplified with the “338” forward primer (5′-ACTCCTACGGGAGGCAGCA-3′) and the “806” reverse primer (5′-GGACTACHVGGGTWTCTAAT-3′). The PCR settings were denaturation at 95 °C for 5 min; 25 cycles of 95 °C for 30 s, 50 °C for 30 s, and 72 °C for 40 s; and final extension at 72 °C for 7 min. To purify the successful PCR products, VAHTSTM DNA magnetic beads was used to remove primers, dimers, salts, and deoxynucleoside triphosphates (dNTPs). Equally molar DNA libraries were prepared for pair-end sequencing using Illumina NovaSeq 6000 (Illumina, San Diego) platform.

### 2.4 Bioinformatics

Raw sequence data were primarily filtered by Trimmomatic (version 0.33) based on the quality of a single nucleotide (Bolger et al., 2014). Identification and removal of primer sequences were performed by Cutadapt (version 1.9.1) (Martin, 2011). Reads were demultiplexed, and paired-end reads were merged using USEARCH (version 10) (Edgar, 2010) followed by chimera removal using UCHIME (version 8.1) (Edgar et al., 2011). High-quality reads were obtained after removing chimeras. To analyze the microbial diversity information of the samples, clean tags were grouped at a 97% sequence similarity using USEARCH in QIIME. Different operational taxonomic units (OTUs) were obtained; then classified and annotated using the SILVA (bacteria) databases.

### 2.5 Data analyses

To compare the prey composition in spider guts, and to investigate how different environmental variables (local field-scale and landscape scale) influence the gut microbial assemblages, all statistical analyses was conducted in R software (version 3.6.3). Differential abundance analyses was carried out at order level using R package ‘DESeq2’ which allows identification of differentially abundant taxa between local field-scale groups: pesticide use (conventional versus organic) and crop identity (Chinese cabbage versus cauliflower). Multivariate orthogonal partial least squares discriminant analyses (OPLS-DA) was performed on the normalized log-transformed OTU abundance of bacterial orders considering as factor the combination of local field-scale groups. OPLS-DA was carried out with the R package ‘rolps’. The p-values and value of fold changes of differentially abundant taxa, and results of OPLS-DA were plotted together.

A distance-based redundancy analyis (dbRDA) model, based on Euclidean distances, was used to unveil the relationships between gut microbes of spiders with both local field-scale groups and landscape gradients. The community matrices were Hellinger transformed before performing the dbRDA. This transformation is often used in zero and one inflated community datasets because it downweighs variables with low counts and many zeros. To test the collinearity among each of the dbRDA model predictors, we used variation inflation factors (VIFs) method (James et al., 2013). Because environmental factors with VIF > 10 had collinearity with other environmental variables, they did not significantly contribute to the model’s variance and were excluded from our final model. Additionally, an ANOVA-like permutation (999) test was used to determine the significance of dbRDA models and each environmental constraint (Legendre et al., 2011).

## 3 Results

### 3.1 Stability of gut bacterial community composition between pesticide uses, crop identity and seasons

At a 97 percent sequence alignment cutoff rate, Illumina Hiseq2500 sequencing generated a total of 15,911,790 (Bacteria-16S:V3+V4) clean paired reads, classified into 1,552 bacterial OTUs. A total of 27 phyla, 63 classes, 148 orders, 282 families, and 589 genera were identified. Among the gut bacterial communities, the dominant phylum was Proteobacteria (45.25%), followed by Firmicutes (24.21%), Actinobacteria (8.74%), Cyanobacteria (7.04%) and Bacteroidetes (5.23%) (Fig. 2). Gammaproteobacteria (38.59%) was the most abundant class, followed by Bacilli (13.55%) and Clostridia (9.25%). Similarly, Enterobacteriales (29.68%) has the highest representation in the overall gut bacterial community composition at the order level, followed by Lactobacillales (11.28%) and Clostridiales (9.25%). In the phylum Proteobacteria, Gammaproteobacteria and Enterobacteriales were the most abundant class and order, respectively. Similarly, Bacilli and Lactobacillales were the dominant order and class, respectively in the phylum Firmicutes (Fig. 2).

**Fig. 2.**
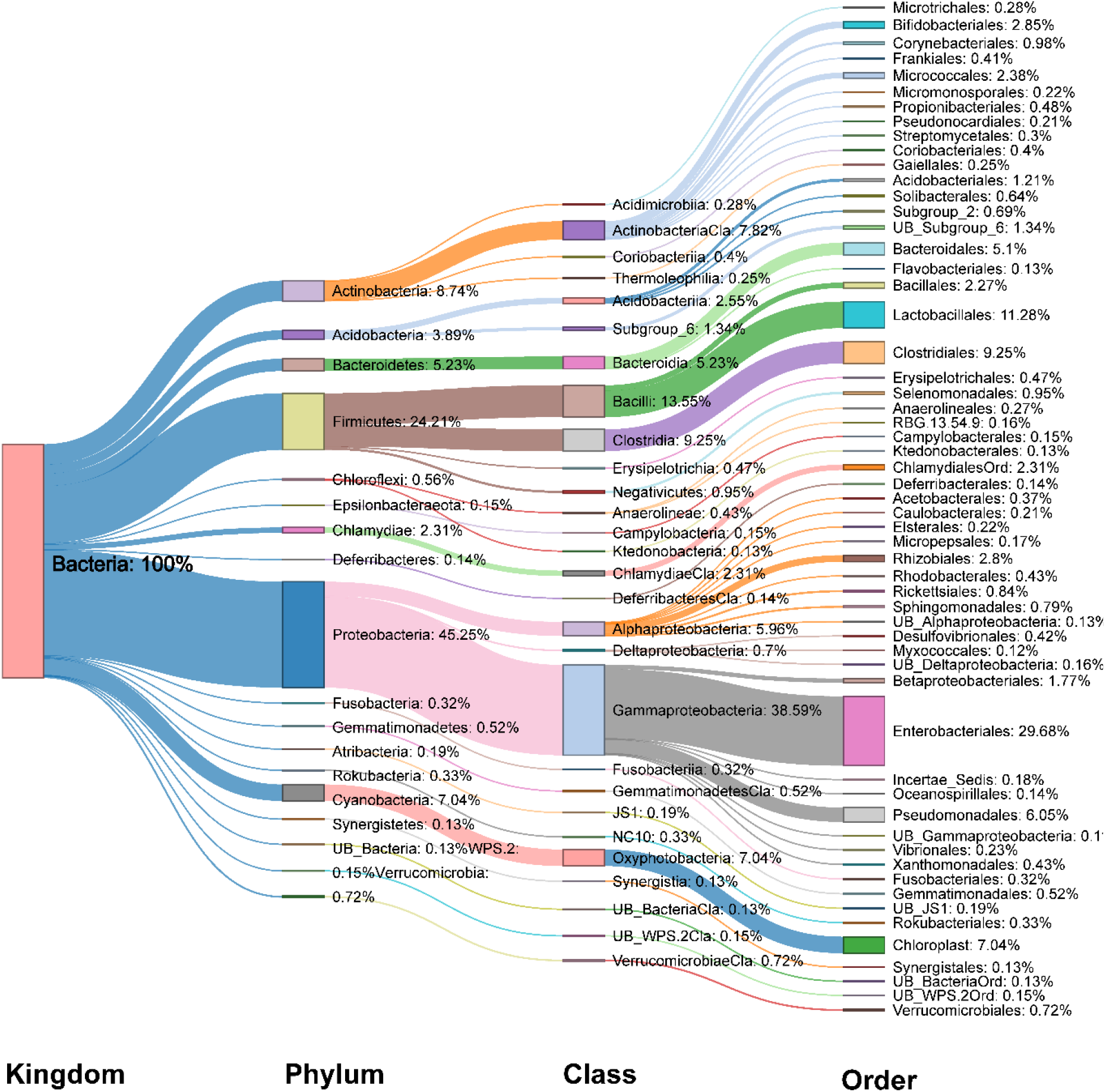
Sankey diagram of relative abundance (%) of dominant bacterial taxa (relative abundance ≥ 0.1%) detected in the gut of spiders.

From the perspective of overall community composition, there was strong similarity across seasons, different pesticide practices, and crop identities. Microbiomes were dominated by Proteobacteria and Firmicutes phyla of bacteria, followed by Actinobacteria, Cyanobacteria, Bacteroidetes and Acidobacteriia (Fig. 3a). At the class level, Gammaproteobacteria and Bacilli represent the highest proportion of bacteria, followed by Clostridia, Actinobacteria, Oxyphotobacteria and Alphaproteobacteria (Fig. 3b). Furthermore, there were no changes to the dominance of the Enterobacteriales, Lactobacillales, Clostridiales, Chloroplast, Pseudomonadales and Bacteroidales orders (Fig. 3c). Similarly, the Enterobacteriaceae, Chloroplast, Lactobacillaceae, Pseudomonadaceae, Lachnospiraceae, Leuconostocaceae and Ruminococcaceae families were dominant regardless of the different farming systems, crop types and seasons (Fig. 3d).

**Fig. 3.**
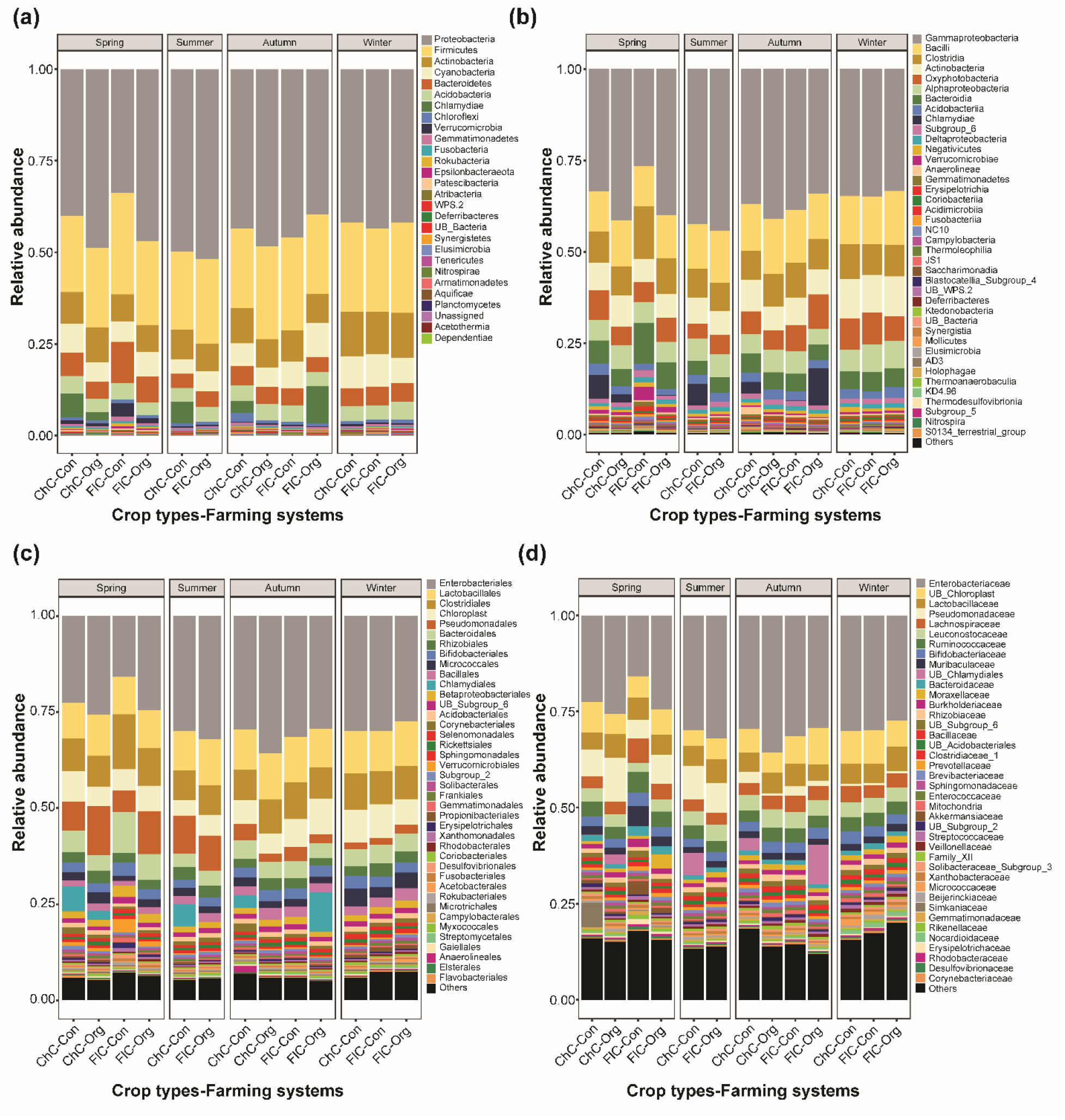
Relative abundance (%) of bacterial **(a)** phylum, top 30 **(b)** classes, **(c)** orders and **(d)** families detected in the gut of spiders. Spiders were captured from different brassica crop type (Chinese cabbage vs cauliflower) fields managed under different farming systems (conventional vs organic) across four seasons. Here, “ChC”, “FlC”, “Con” and “Org” represent Chinese cabbage, cauliflower, conventional and organic respectively.

### 3.2 Effects of pesticide uses and crop identity on gut bacterial community composition

A differential abundance analyses and OPLS-DA was performed to identify the bacterial orders that contributed the most to the variance in the gut of spiders collected from different pesticide uses and crop identities in different season. The overall gut bacterial composition was clearly different between different pesticide uses and crop identities at all four seasons (see inserted boxes of Figs. 4, S1-S3).

**Fig. 4.**
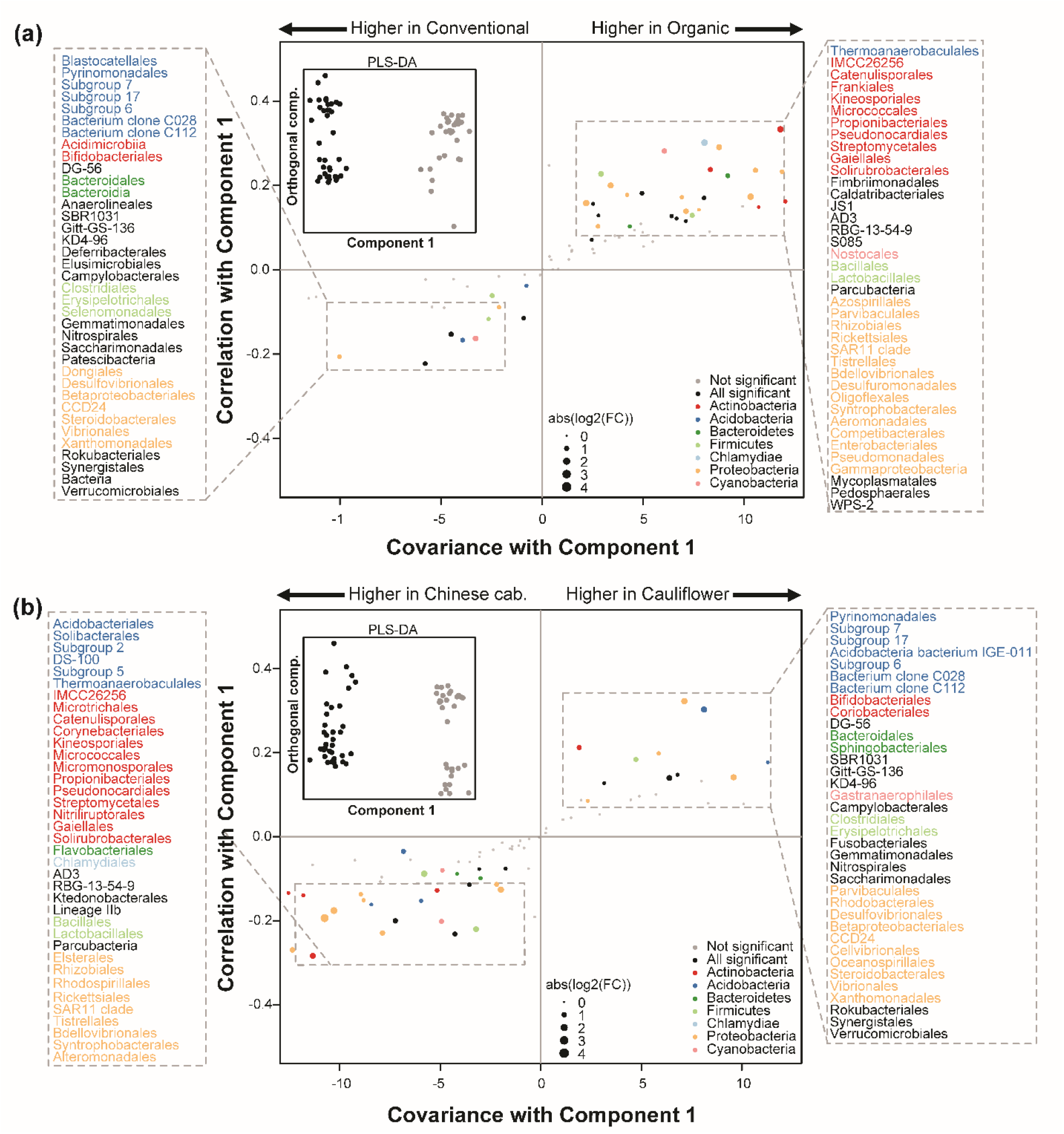
Gut microbiome changes of spiders between **(a)** pesticide uses (conventional versus organic) and **(b)** crop identity (Chinese cabbage versus cauliflower) in spring. The inside black-line boxes show the orthogonal partial least squares discriminant analyses (OPLS-DA) performed on the relative abundance of 148 bacterial orders. S-plots was generated in the main boxes from the results of OPLS-DA and differential abundance analyses. Each point shows the covariance (x-axis) and correlation (y-axis) from the predictive components of OPLS-DA model. Size of each point shows the value of fold change (FC) obtained from differential abundance analyses. Orders that were not robustly significantly (padj > 0.01) different between pesticide uses and crop identities are plotted in grey. Significant (padj < 0.01) families belonging to the top phyla (overall relative abundance > 2%) are plotted in color and significant (padj < 0.01) families belonging to other phyla (overall relative abundance < 2%) are plotted in black. padj corresponds to the *p*-value adjusted for multiple correlation testing using the Benjamini–Hochberg method.

A discriminant analyses on spider samples collected from the different pesticide uses and different crop identities revealed strongest differences in the relative abundance of gut bacterial taxa during spring (Fig. 4) and autumn (Fig. S1). In spring, the 76 gut bacterial orders contributed the most in discriminating the pesticide uses (conventional verses organic) (Fig. 4a), and 73 orders contributed to discriminating the crop identities (Chinese cabbage verses cauliflower) (Fig. 4b). The most of differentially abundant orders between pesticide uses were mainly belong to the Acidobacteria (e.g., Blastocatellales, Pyrinomonadales and Thermoanaerobaculales), Actinobacteria (e.g., Acidimicrobiia, Bifidobacteriales, Catenulisporales, Frankiales, Kineosporiales, Micrococcales, Propionibacteriales, Pseudonocardiales and Streptomycetales), Firmicutes (e.g., Clostridiales, Erysipelotrichales, Selenomonadales, Bacillales and Lactobacillales) and Proteobacteria (e.g., Dongiales, Desulfovibrionales, Betaproteobacteriales, Steroidobacterales, Azospirillales, Parvibaculales, Rhizobiales, Enterobacteriales, Pseudomonadales and Gammaproteobacteria) phyla (Fig. 4a). Similarly, during the spring season, the differences of gut bacterial orders concerning the crop identities were also exclusively assigned to the Acidobacteria (e.g., Acidobacteriales, Solibacterales, Thermoanaerobaculales and Pyrinomonadales), Actinobacteria (e.g., Microtrichales, Catenulisporales, Corynebacteriales, Kineosporiales, Micrococcales, Pseudonocardiales, Bifidobacteriales and Coriobacteriales), Firmicutes (e.g., Bacillales, Lactobacillales, Clostridiales and Erysipelotrichales) and Proteobacteria (e.g., Elsterales, Rhizobiales, Rhodospirillales, Rickettsiales, Tistrellales, Parvibaculales, Rhodobacterales, Desulfovibrionales and Betaproteobacteriales) phyla (Fig. 4b).

In the autumn, discriminant analyses identified several significantly differentially abundant taxa in the gut of spiders collected from different pesticide uses and crop identities. The abundance of the 26 orders were significantly different between pesticide uses (organic verses conventional) fields (Fig. S1a), and 31 bacterial orders contributed to the differentiation between crop identities (Chinese cabbage verses cauliflower) (Fig. S1b). Differences concerned between pesticide uses almost exclusively assigned to the Acidobacteria (e.g., Blastocatellales, Subgroup 7 and Thermoanaerobaculales), Actinobacteria (e.g., Kineosporiales, Pseudonocardiales and Streptomycetales) and Proteobacteria (e.g., Acetobacterales, Caulobacterales, Rhodobacterales, Enterobacteriales and Alphaproteobacteria) phyla (Fig. S1a). The spider collected from Chinese cabbage had the significantly higher abundance of phyla Acidobacteria (e.g., Subgroup 7 and soil Bacterium clone C028), Actinobacteria (e.g., Corynebacteriales and Pseudonocardiales), Bacteroidetes (Bacteroidales, Chitinophagales and Sphingobacteriales) and Proteobacteria (e.g., Dongiales, SAR11 clade and Vibrionales) (Fig. S1b). On the other hand, the guts of spider collected from cauliflower was significantly differentially abundant with phyla Actinobacteria (e.g., Frankiales, Kineosporiales and Micromonosporales) and Proteobacteria (e.g., Azospirillales, Elsterales and Rickettsiales) (Fig. S1b).

In winter, a total of 29 bacterial orders (e.g., Acidobacteriales, Solibacterales, Pyrinomonadales, Catenulisporales, Cytophagales, Bacillales, Caulobacterales and Elsterales) were significantly differentially abundant in organic fields compared with the gut of spiders collected form conventionally managed fields were significantly differentially dominated with 14 bacterial orders (e.g., Kineosporiales, Chloroplast, Rickettsiales and Bdellovibrionales) (Fig. S2a). Moreover, 13 bacterial orders (e.g., Corynebacteriales, Frankiales, Actinobacteria and Sphingomonadales, Desulfuromonadales and Myxococcales) were significantly differentially dominant in the guts of spiders collected from cauliflower fields compared with samples collected from Chinese cabbage had 12 significantly differentially abundant bacterial orders (e.g., Bifidobacteriales, Coriobacteriales, Sphingobacteriales, Clostridiales and Micropepsales) (Fig. S2b). In summer, however, the total of 20 gut bacterial orders (e.g., Aminicenantales, Streptosporangiales, Chloroplast, Lactobacillales, Acetobacterales and Rickettsiales) were significantly higher in spiders collected from cauliflower fields under organic management practices, whilst only five gut bacterial orders (e.g., Acetothermiia, Propionibacteriales, Sphingobacteriales and Oligoflexales) were significantly differentially abundant in spiders captured from Chinese cabbage field under conventional management practices (Fig. S3). These results of discriminant analyses highlighted that spider showed a clear divergence of certain gut microbes between different pesticide uses and crop identities across different seasons.

### 3.3 Local scale groups and landscape gradients as determinants of gut microbial assemblages

The abundance and OTUs Shannon-diversity analyses revealed effects of landscape features, seasons, crop types and farming systems whether considering of the entire gut microbiome of spiders or a subset of the most numerous nine orders (Fig. 5). The differences in the abundance and OTUs Shannon-diversity of microbes detected in the gut of spiders were significant between local- and landscape-factors across different seasons (dbRDA model permutation test. Only the “spatial scales (i.e., differing sized radii around the focal field)” was found to be redundant (having VIF>10) in the final dbRDA models of both abundance and Shannon-diversity of gut microbes, therefore it was removed. The first two axes in the final dbRDA model explained 96% of total variability in the assemblage structure of gut microbes based on their abundance and 95% based on their OUT’s Shannon-diversity. Consistent with the PERMANOVA results, the permutation of dbRDA also showed that all the local and landscape factors significantly explained the variability observed in the assemblage structure of both abundance and Shannon-diversity of the gut microbes in spiders.

**Fig. 5.**
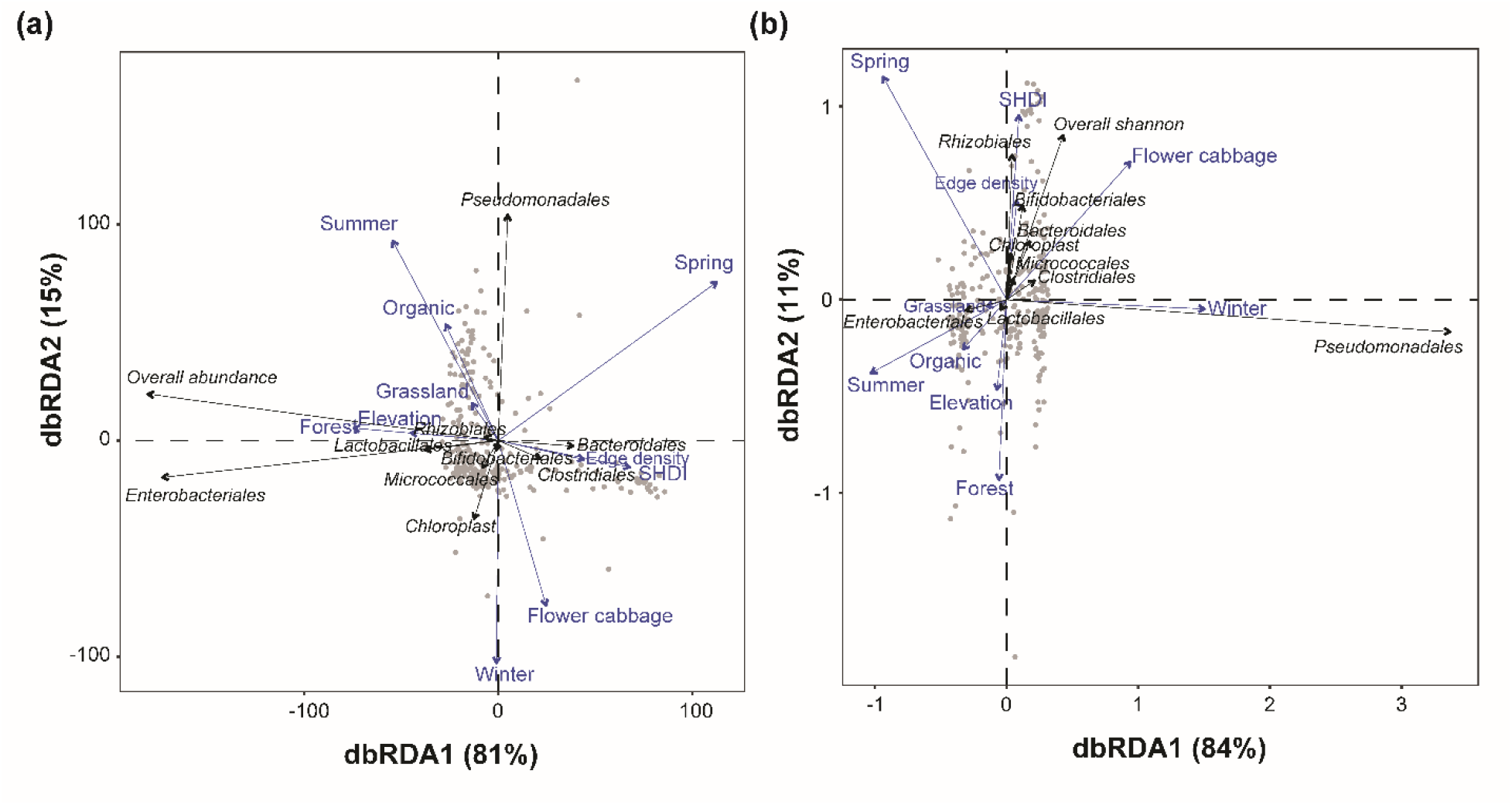
Distance based redundancy analyses (dbRDA) illustrating the associations of overall and top ten bacterial taxa with various environmental factors in terms of **(a)** abundance and **(b)** diversity in the gut of spiders. For each variable, the arrows’ length and orientation indicate the magnitude of explained variance. The association between prey taxa detected in spider guts and explanatory factors represented by the perpendicular distance between them (below-90°= positive association and above-90° = negative association). The association is larger when the perpendicular distance is less.

The higher proportions of forests, grasslands and edge density accounted for significantly higher fractions of the variability in assemblages of gut microbes in terms of both abundance and Shannon-diversity (Table 1). The compositional diversity of landscape (SHDI), however, accounted for significantly explaining the higher fractions of the variability in the assemblages of gut microbes in terms of only their OTUs Shannon-diversity (OTUs Shannon-diversity – *F* = 2.000, *p* = 0.043). At local field scale, both abundance and OTUs Shannon-diversity of gut microbes were significantly influenced by the crop identity, pesticide use, seasons, and elevation gradients. The seasons and crop identity, on the other hand, accounted for explaining the higher fractions of the variability in assemblages of gut microbes in term of both abundance and Shannon-diversity (Table 1).

**Table 1.** ANOVA table to test the significance of each predictor in RDA model explaining the variance of bacterial communities in the gut of spiders.

| Variables | Abundance |  | Diversity |  |
| --- | --- | --- | --- | --- |
|  | F | P-value | F | P-value |
| RDA Models | 40.012 | 0.001 | 58.415 | 0.001 |
| *Land use variables* |  |  |  |  |
| Edge density | 15.108 | 0.001 | 7.808 | 0.001 |
| SHDI | 1.064 | 0.326 | 2.000 | 0.043 |
| Forest | 25.961 | 0.001 | 7.926 | 0.001 |
| Grassland | 3.706 | 0.02 | 4.260 | 0.006 |
| *Other variables* |  |  |  |  |
| Crop identity | 47.005 | 0.001 | 58.740 | 0.001 |
| Pesticide use | 19.935 | 0.001 | 5.979 | 0.001 |
| Season | 84.635 | 0.001 | 162.692 | 0.001 |
| Elevation | 33.436 | 0.001 | 9.358 | 0.001 |

The overall gut microbial abundance and the abundances of Enterobacteriales, Lactobacillales, Rhizobiales were higher in fields surrounded with high proportion of forest and grassland located at higher elevation during autumn season. Conversely, the abundance of Bacteroidales and Clostridiales gut microbes was strongly associated with higher edge density and higher SHDI. Cauliflower (as the focal crop type) fields had higher abundance of Bifidobacteriales, Micrococcales and Chloroplast microbes during winter (Fig. 5a). For the OTUs Shannon-diversity of gut microbial taxa, the overall Shannon-diversity and the Shannon-diversity of Rhizobiales, Bifidobacteriales, Bacteroidales, Chloroplast, Clostridiales and Micrococcales were strongly associated with the cauliflower (as the focal crop type), and the higher edge density as well as the higher landscape compositional diversity (SHDI) during spring season. The Shannon-diversity of Pseudomonadales in the gut of spiders was higher during winter season. Conversely, the Shannon-diversity of Lactobacillales and Enterobacteriales in the gut of spiders was strongly belonging to the organic fields with higher proportions of grassland and forest patches, and higher elevation during summer season (Fig. 5b).

## 4 Discussion

Anthropogenic changes in agroecosystems are inducing arthropod adaptation to ecological disturbances at an unprecedented scale (Gurr et al., 2016; Karp et al., 2018; Michalko et al., 2019; Pearson et al., 2014; Saqib et al., 2022, 2020; Thomas et al., 2004; Tuck et al., 2014). Gut microbes strongly influence the host fitness, including development, reproduction, host nutrition, stress tolerance against various ecological disturbances (Engel and Moran, 2013; Jang and Kikuchi, 2020). We have demonstrated that variations in different local environmental conditions and composition of surrounding landscape distinctly altered the gut microbiota of spiders. This study provides powerful insight into the plasticity of bacterial diversity and abundance in the gut of the most abundant predator in agricultural crops and demonstrates for the first time the effects of seasons, crop identity, pesticide use, and surrounding landscape structure on the gut microbiota. In this study, we detected 27 phyla, 64 classes, 148 orders, 282 families and 589 genera of bacteria. The dominant bacterial taxa (such as Proteobacteria, Firmicutes, Actinobacteria, Cyanobacteria, Bacteroidetes and Acidobacteria) represents more than 80% of the entire gut bacterial community of spiders. We have demonstrated that the relative abundance of several gut bacterial taxa of spiders were distinctly influenced by different local field-scale and landscape-scale variables.

Prior studies demonstrated that composition of diet is closely related to gut microbiota of arthropods (Dong et al., 2018; Gupta and Thorsteinson, 1960; Yang et al., 2020). In our study, the composition of the gut microbiota was substantially discriminated by crop identity. Spiders are known as generalist predators and prey on a diverse range of herbivores as well as on other predator species (intraguild predation) (Hambäck et al., 2021; Saqib et al., 2021). The availability and feeding on different kinds of crops acts as a vital source for diverse gut microbial population for plant-eating insects. Feeding on different crop types cause differences in herbivores gut microenvironment, which may lead to the variation in gut microbial diversity of their associated predators. For example, the gut bacteria community structure of *Monochamus alternatus*, *Psacothea hilaris* (Coleoprera: Cerambycidae), *Rhodococcus* and *Achromobacter* genera have a strong link with natural diet samples, and the *Buttiauxella* and *Kluyvera* genera showed a significant correlation with artificial diet samples (Kim et al., 2017). Wild crickets, *Teleogryllus oceanicus* (Orthoptera: Gryllidae) have five bacterial phyla (*Cyanobacteria*, *Fusobacteria*, *Lentisphaerae*, *Planctomycetes* and *Synergistetes*) that were not detected in lab-reared crickets (Ng et al., 2018). In addition, some Lepidopteran insects such as *Thaumetopoea pityocampa* and *Plutella xylostella* also showed an interplay of strong interactions between the insect gut bacterial community and host plants (Strano et al., 2018). The abundance of Proteobacteria phylum was also found to be less in the larvae feeding on *Pinus halepensis* (Pinales: Pinaceae) than those collected on *Pinus nigra* subsp. *laricio* or *Pinus pinaster* (Strano et al., 2018). The underlined above findings suggested that generalist predators foraging on diverse prey range (including herbivores as well as intraguild prey) which feed on different crop plants will eventually determine the assemblage structure of predator’s gut microbiota.

Similar to the impacts of the host crops, there were considerable effects of different pesticide uses on discriminating the assemblages of gut microbial taxa in spiders. It is widely reported that increasing organic farming would lead to the higher farmland biodiversity (Batáry et al., 2011; Fuller et al., 2005; Garratt et al., 2011; Schmidt et al., 2005). Plant species density has been reported to be higher in organic fields (both inside the crop fields and adjacent non-crop areas) than in conventional ones (Chateil et al., 2013), which may provide a range of resources, and thus contribute to the more diverse assemblages of arthropods. Conventional farms receive more synthetic herbicides to ensure weed control, resulting in lower plant species diversity (both within the crop fields as well as in adjacent areas), which ultimately has severe impacts on the diversity of inhabiting arthropods. Similarly, study have reported the variations in chemical composition of soil as well as plant metabolites (e.g., pH, organic matter, nutrients, root exudates) between conventional and organic farming systems (Armalytė et al., 2019; Cubero-Leon et al., 2018). These changes of plant metabolites may also drive changes in metabolic structure of herbivores (Kešnerová et al., 2017; Yang et al., 2020), which might eventually influence the assemblages of gut microbiota of herbivores and their associated predators.

Furthermore, continuous application of pesticide has developed insecticide resistance in pests (Brattsten et al., 1986). Especially, neonicotinoids are neurotoxic systemic insecticides widely employed in plants to suppress sucking insect-pests (Nauen and Denholm, 2005). They may affect both target as well as beneficial insects. Gut microbiota of insects seems to be involved in pesticide resistance and biodegradation of Neonicotinoids. Few researchers examined how insect gut microbial populations affect pesticide resistance. Kikuchi et al. (2012) showed that the gut symbiont Burkholderia mediates pesticide resistance in *Riptortus pedestris* (Hemiptera) and that this resistance may be horizontally transferred to other insects. Direct biodegradation of pesticides by gut microbiota, such as *R. pedestris* (Kikuchi et al., 2012) and *B. dorsalis* (Cheng et al., 2017), and immune modulation, in which microbiota induce the development of an innate immune response (Broderick et al., 2010), have been proposed as drivers of resistance development. Bacteria including *Bacillus aerophilus, Klebsiella pneumonia*, Mycobacterium sp, Pseudomonas sp. 1G, Pseudoxanthomonas, Stenotrophomonas sp., Rhodococcus sp. found to be involved in pesticide resistance and biodegradation of neonicotinoids (Xia et al., 2018). In our study most of these bacteria found in the gut of spiders. However, detailed studies are required to unveil the whole underlying mechanisms of pesticide resistance in natural enemies and to explore the role different genes and enzyme of gut microbes involved in the biodegradation of pesticides.

A study conducted on cricket, *Gryllus veletis* revealed that, the insect gut microbiota changed with season, and overwintering affects the composition of its gut microbiome compared to spring field *G. veletis*; where the relative abundance of Proteobacteria increased in the gut (Ferguson et al., 2018). Pseudomonadales (belong to phylum Proteobacteria) were found to be dominant in spiders sampled during summer, whereas Lactobacillales (phylum Firmicutes) and Enterobacteriales (phylum Proteobacteria) were dominant in the bacterial profile of samples collected during autumn. In another study, Proteobacteria exhibited a significant variation with the seasons in culex (Duguma et al., 2017), indicating that the Proteobacteria may be associated with seasonal changes in factors such as temperature. Overall, our results showed that spider samples collected in summer had the lowest assemblages in terms gut microbiocidal diversity compared to samples collected in spring, autumn and winter. This could be explained by the fact that in the summer, few farmers grow brassica vegetables because the summer temperature can reach up to 40 ℃ and above in Fuzhou province. Research shows that high temperature could affect the growth of plants, and the growth and development of insects, even alter the insect’s feeding behavior, and indirectly can influence the assemblages of gut microbiota of arthropods (Sepulveda and Moeller, 2020).

To date, surrounding habitat and the ecological conditions shaping the microbial community in the gut of generalist predators has received comparatively little attention. In our study, we showed that all the landscape variables significantly linked to the variations observed in assemblages of gut microbiota in spiders; particularly varying proportions landscape composition diversity (SHDI), forest patches, grassland patches and edge density were the factors which showed significant influences. In several studies it has been reported that varying proportions of different land uses and edge density (Gallé et al., 2018; Landis et al., 2000; Perović et al., 2015) as well as the landscape compositional diversity (Zhang et al., 2021) in the surrounding landscape may alter the assemblage patterns of farmland biodiversity, which may likely also have significant impact on driving the distribution of microbes at various trophic levels within an ecosystem. In a characterization study of several insect species and their associated gut microbiota, the relative occurrences of microbes were found to vary according to the surrounding environmental habitats of the insects (Yun et al., 2014). Overall, these results suggest that the surrounding environment that spiders are exposed to, including varying proportions of different land uses, environmental conditions, or prey range, may highly affect their gut microbial assemblages.

## 5 Conclusions

In summary, the results of our study provide the evidence that the varying environmental exposures at local and broad landscape levels had distinct impacts on the assemblages of gut microbiota of spiders. In addition, our results underscore the possibility that key environmental factors may influence the functionally important gut microbes (such as Proteobacteria and Firmicutes), and that future laboratory-based studies are imperative for understanding their functional role in growth, development, nutrition and reproduction of spiders. Although it is hard to determine the spatial origin of microbes inside the body of generalist predators, it is paramount to assess or establish whether these microbes were directly acquired from the environment or a result of an indirect acquisition through predation on varying prey species or vertical transmission from the mother spider. This study, however, highlights the possible complex interplay between host, its gut microbial community, and the key environmental factors (at various local- and landscape-scales), and identifies the key bacteria taxa to target in future investigations.

## Acknowledgments

This work was financially supported by the Natural Science Foundation of Fujian Province, China (2022J06013), the National Natural Science Foundation of China (No. 31972271), State Key Laboratory of Ecological Pest Control for Fujian and Taiwan Crops, Joint International Research Laboratory of Ecological Pest Control, Fujian-Taiwan Joint Innovation Centre for Ecological Control of Crop Pests, International science and technology cooperation and exchange program of FAFU (KXb16014A), and the Thousand Talents Program and the “111” Program in China.

## Conflict of interest

The authors declare no conflict of interest.

## Author contributions

H.S.A.S., M.Y., S.Y. and G.M.G. conceived and designed the experiments. H.S.A.S., K.S.A. and L.S. conducted field sampling and lab experiments. H.S.A.S. performed the data analyses, prepared figure and tables. H.S.A.S., G.P., M.U.G and G.M.G. interpreted the data and wrote the paper. P.L., S.Y. and G.P. assisted in the data analyses. All authors revised the final version and gave their approval for submission.

## Data availability

All data supporting the results of this article are included in the published article and its supplementary information files.

## Notes

### Competing Interest Statement

The authors have declared no competing interest.

