## Supplementary material for "Gut microbiota assemblages of generalist predators are driven by local- and landscape-scale factors": Fig. S1-3

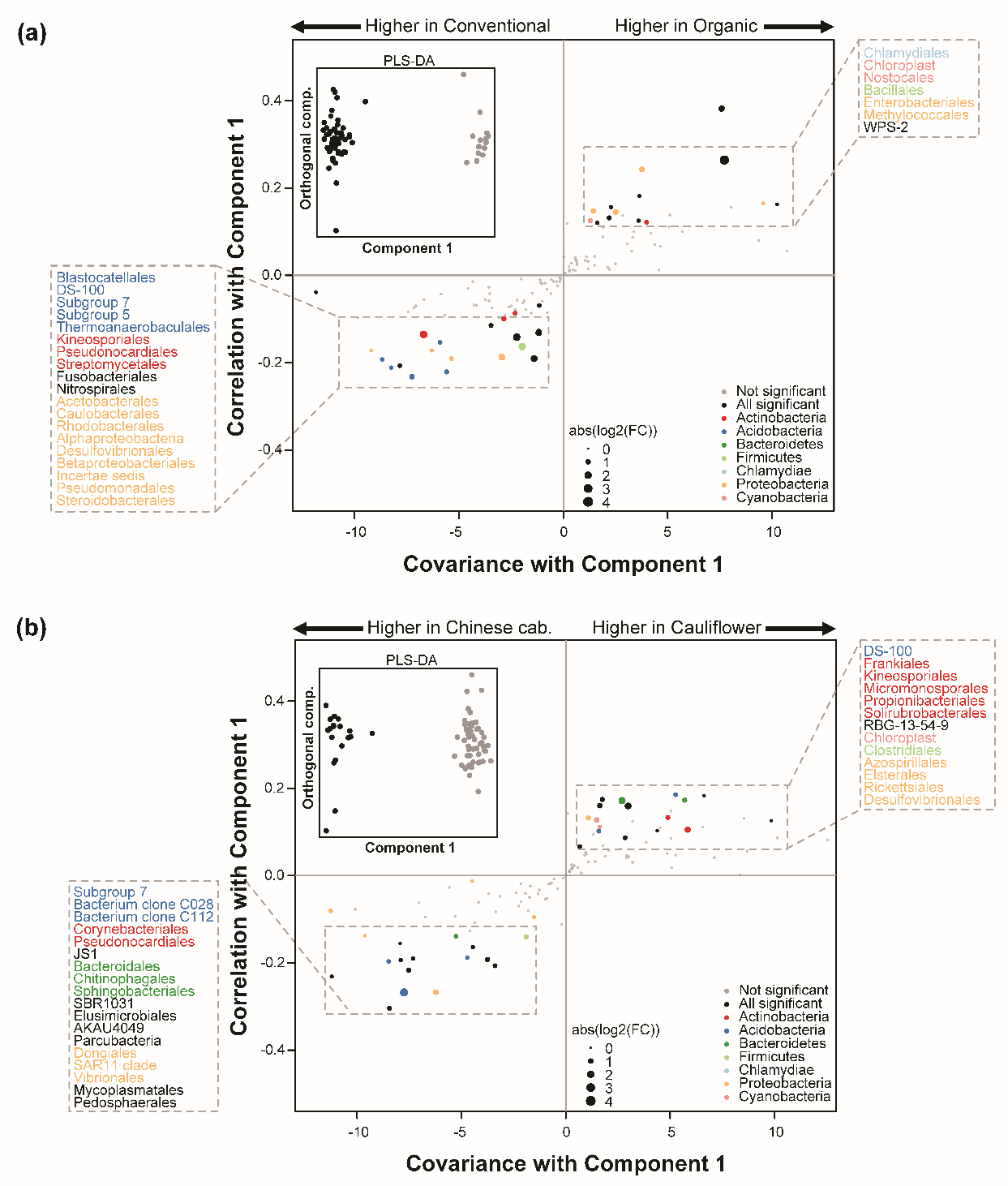

**Fig. S1** Gut microbiome changes of spiders between **(a)** pesticide uses (conventional versus organic) and **(b)** crop identity (Chinese cabbage versus cauliflower) in autumn. The inside boxes black-line show the orthogonal partial least squares discriminant analyses (OPLS-DA) performed on the relative abundance of 148 bacterial orders. S-plots was generated in the main boxes from the results of OPLS-DA and differential abundance analyses. Each point shows the covariance (x-axis) and correlation (y-axis) from the predictive components of OPLS-DA model. Size of each point shows the value of fold change (FC) obtained from differential abundance analyses. Orders that were not robustly significantly (padj > 0.01) different between pesticide uses and crop identities are plotted in grey. Significant (padj < 0.01) families belonging to the top phyla (overall relative abundance > 2%) are plotted in color and significant (padj < 0.01) families belonging to other phyla (overall relative abundance < 2%) are plotted in black. padj corresponds to the *p*-value adjusted for multiple correlation testing using the Benjamini–Hochberg method.

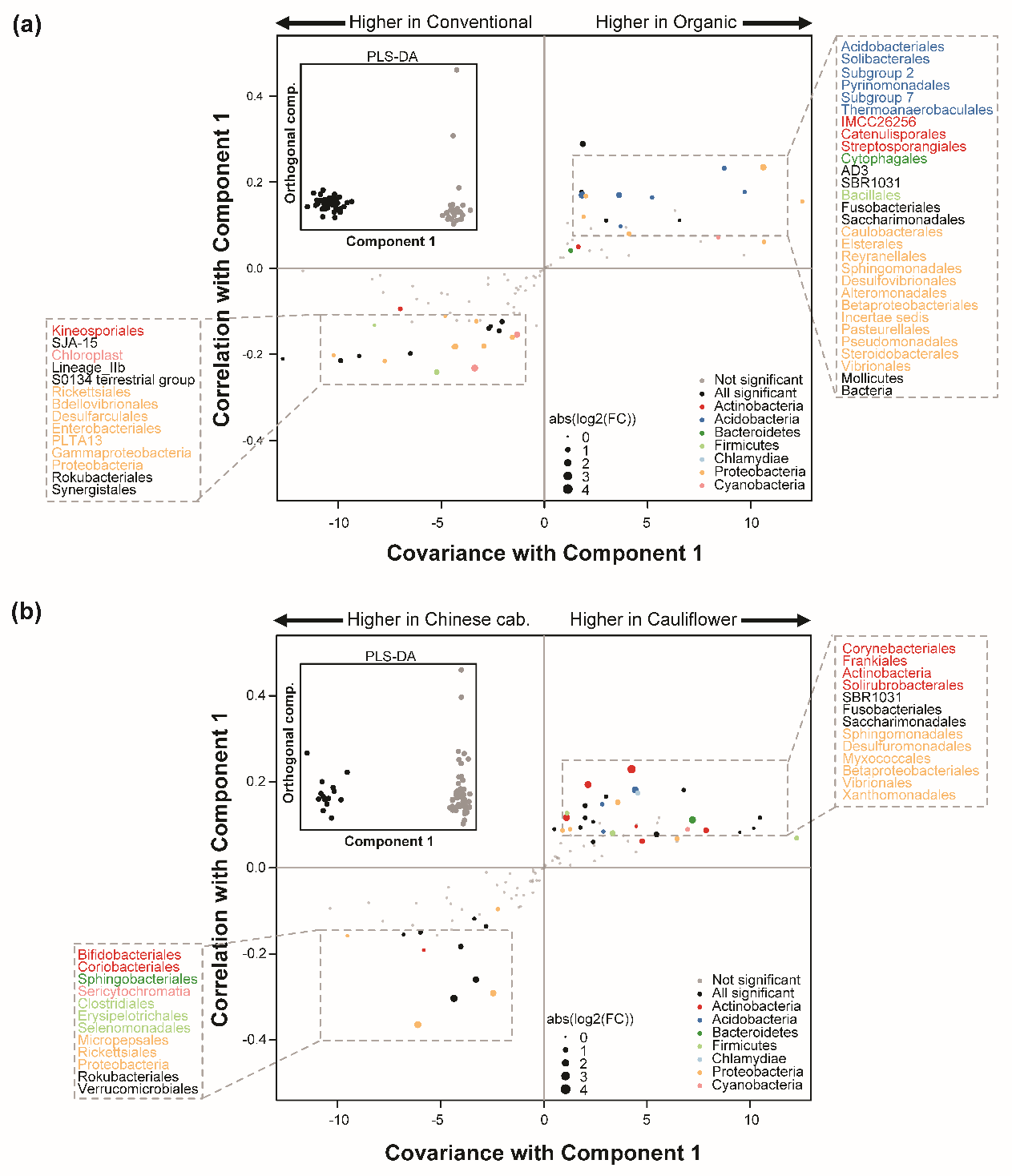

**Fig. S2** Gut microbiome changes of spiders between **(a)** pesticide uses (conventional versus organic) and **(b)** crop identity (Chinese cabbage versus cauliflower) in winter. The inside black-line boxes show the orthogonal partial least squares discriminant analyses (OPLS-DA) performed on the relative abundance of 148 bacterial orders. S-plots was generated in the main boxes from the results of OPLS-DA and differential abundance analyses. Each point shows the covariance (x-axis) and correlation (y-axis) from the predictive components of OPLS-DA model. Size of each point shows the value of fold change (FC) obtained from differential abundance analyses. Orders that were not robustly significantly (padj > 0.01) different between pesticide uses and crop identities are plotted in grey. Significant (padj < 0.01) families belonging to the top phyla (overall relative abundance > 2%) are plotted in color and significant (padj < 0.01) families belonging to other phyla (overall relative abundance < 2%) are plotted in black. padj corresponds to the *p*-value adjusted for multiple correlation testing using the Benjamini–Hochberg method.

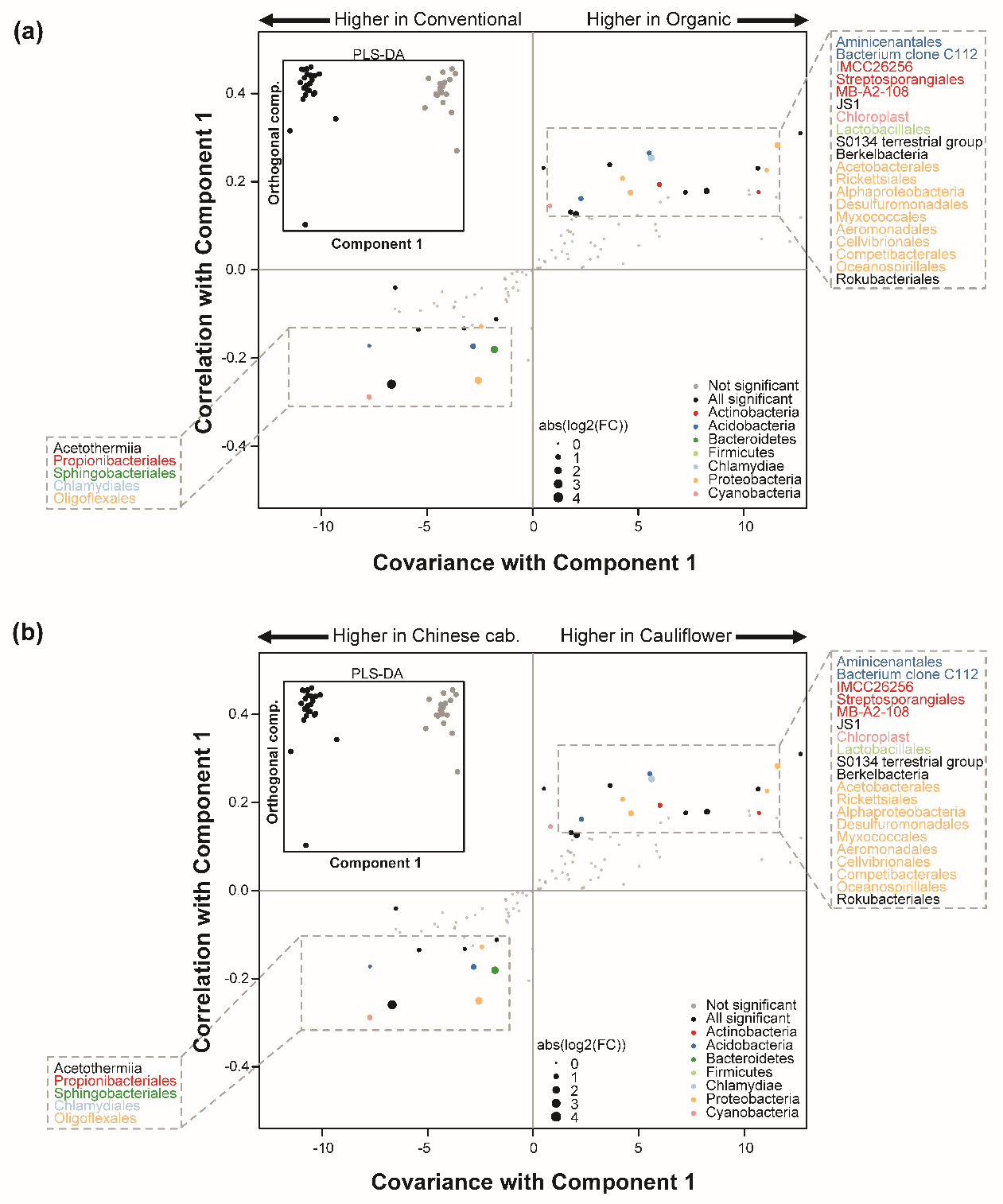

**Fig. S3** Gut microbiome changes of spiders between **(a)** pesticide uses (conventional versus organic) and **(b)** crop identity (Chinese cabbage versus cauliflower) in summer. The inside black-line boxes show the orthogonal partial least squares discriminant analyses (OPLS-DA) performed on the relative abundance of 148 bacterial orders. S-plots was generated in the main boxes from the results of OPLS-DA and differential abundance analyses. Each point shows the covariance (x-axis) and correlation (y-axis) from the predictive components of OPLS-DA model. Size of each point shows the value of fold change (FC) obtained from differential abundance analyses. Orders that were not robustly significantly (padj > 0.01) different between pesticide uses and crop identities are plotted in grey. Significant (padj < 0.01) families belonging to the top phyla (overall relative abundance > 2%) are plotted in color and significant (padj < 0.01) families belonging to other phyla (overall relative abundance < 2%) are plotted in black. padj corresponds to the *p*-value adjusted for multiple correlation testing using the Benjamini–Hochberg method.
